## Supplemental Material for "Single cells can resolve graded stimuli"

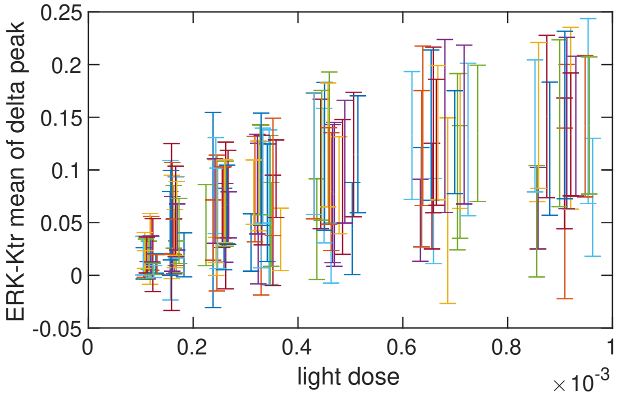


FIG. S1: **Dose-responses of all cells** from one stage position in the main dataset (20 repeats). The bars are centered at the mean ERK response value across the 20 repeats to the measured light doses, and the error bars depict the standard deviation.

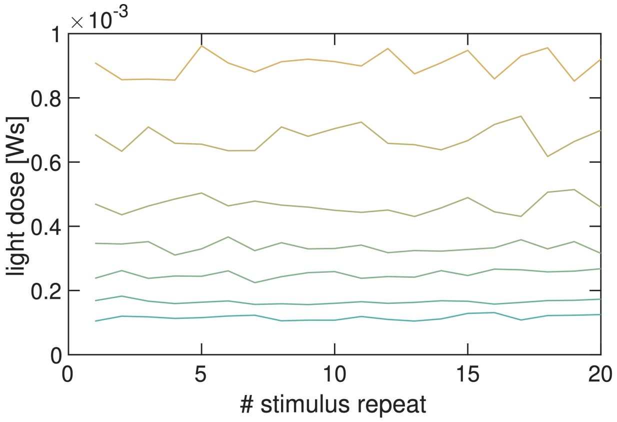


FIG. S2: **The variability between repeats of light doses.** Each of the 7 light doses is represented by a line connecting the 20 repeats of the dose.

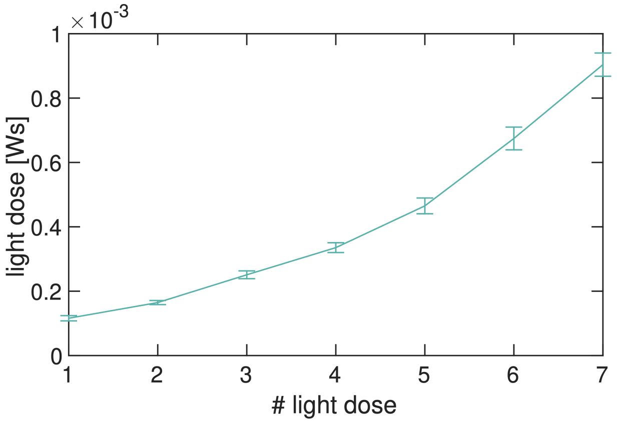


FIG. S3: **The average measured light doses**, connected by a line. The error bars show the standard deviation for the 20 repeats of each light dose.

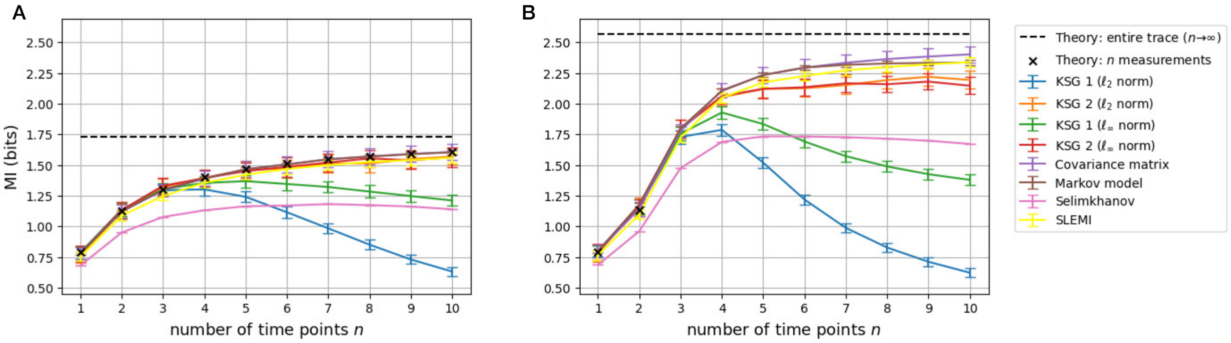


FIG. S4: **Benchmarking of mutual information estimators**, plots from [1] for overdamped (a) and underdamped (b) dynamics, here with the addition of SLEMI algorithm. The plots demonstrate that SLEMI over-performs commonly used estimators. Number of samples N= 20000.

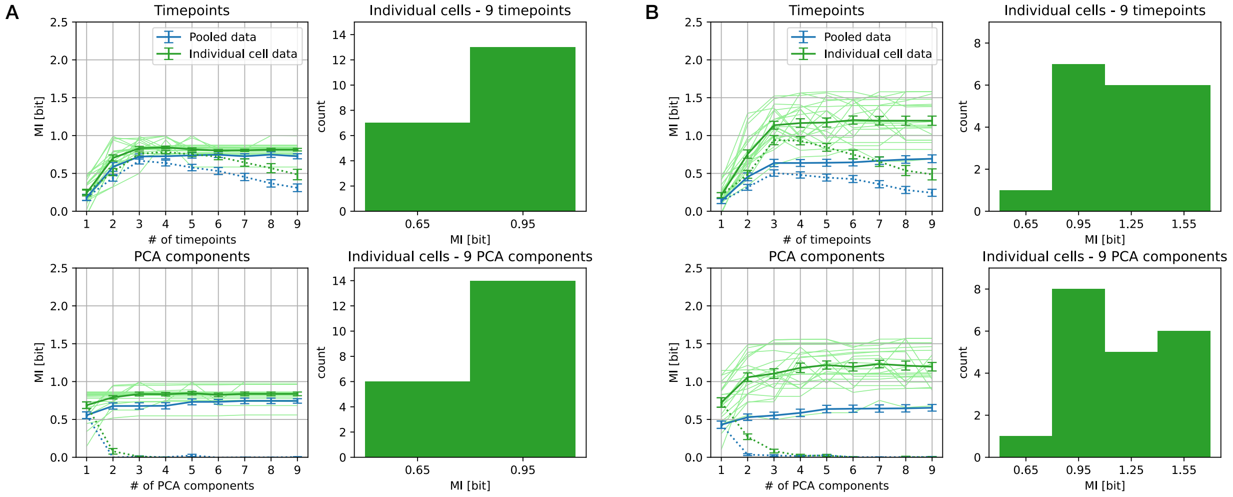


FIG. S5: **Mutual information for different number of input states.** Mutual information estimated for the cells of the main dataset with 20 repeats, in panel (A) taking into account only the first (weakest) and the last (7th, strongest) stimulus, while in (B) the fourth (mid-range) stimulus is additionally taken into account.

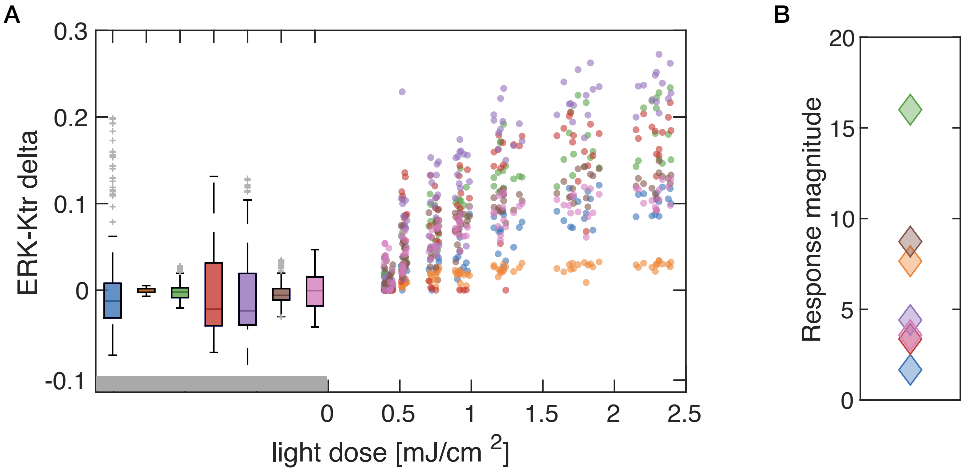


FIG. S6: **Comparison between the endogenous ERK fluctuations and ERK response to light stimuli** for all cells from one stage position of the main dataset with 20 repeats (A). In (B), response magnitude is calculated as the ratio between the maximal response to the stimulus and the standard deviation of the endogenous fluctuations.

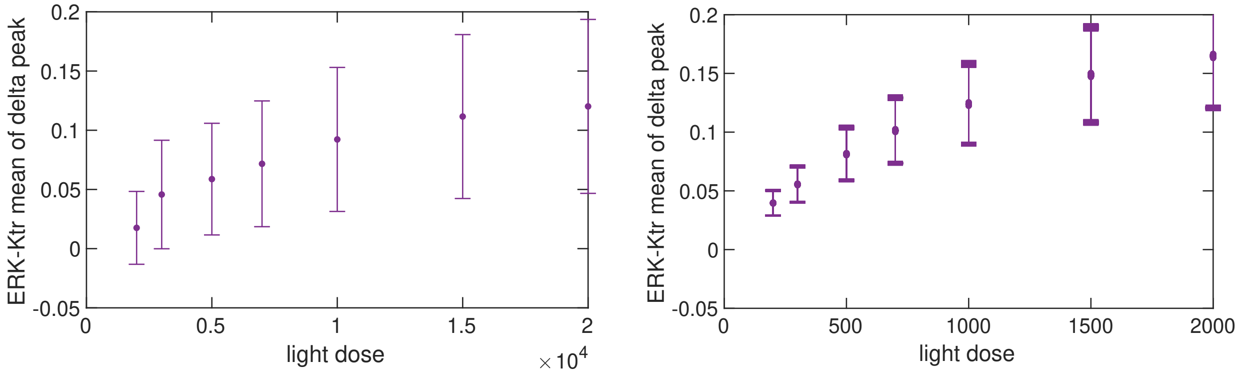


FIG. S7: **Experimental (left) and simulated (right) dose-responses.** The dose-responses in the simulations resemble the experimental ones. Note that the relationship between the simulated light doses corresponds to the experimental setting, not their magnitude displayed the plot.

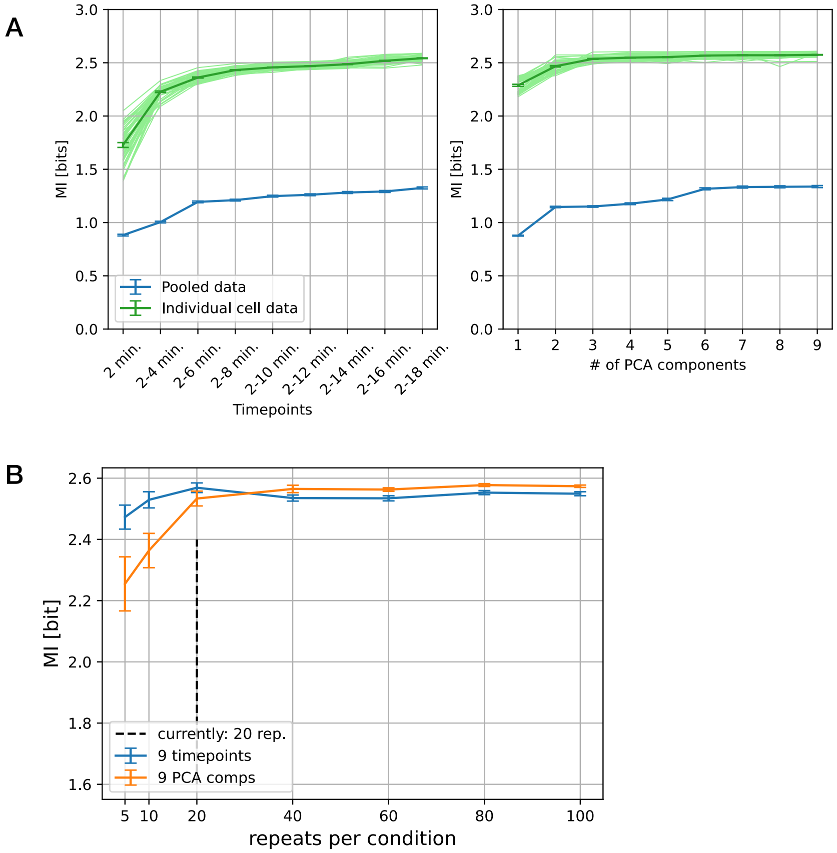


FIG. S8: **Simulations of optogenetic activation of the MAPK cascade in conditions of low noise.** Analogously to the Fig.4 in the main text, the trajectories resulting from the simulations described in the main text were modified by adding 0.0001 Gaussian white noise. **(A)** Mutual information contained in the simulated cells' ERK response trajectories was calculated using the time point feature reduction (left) and PCA (right). The results from pooled cell responses is shown in blue, and the results from single cell responses in green (individual cells in light green, the average of single cell responses in dark green). **(B)** The estimation of mutual information in simulated trajectories for different number of repeats per condition. The information is estimated using the first 9 time points (blue) and PCA (orange). The value of mutual information increases with the increasing sample size, stabilizing at around 20 repeats. The black dashed line is drawn at 20 repeats per condition, the current experimental limit.

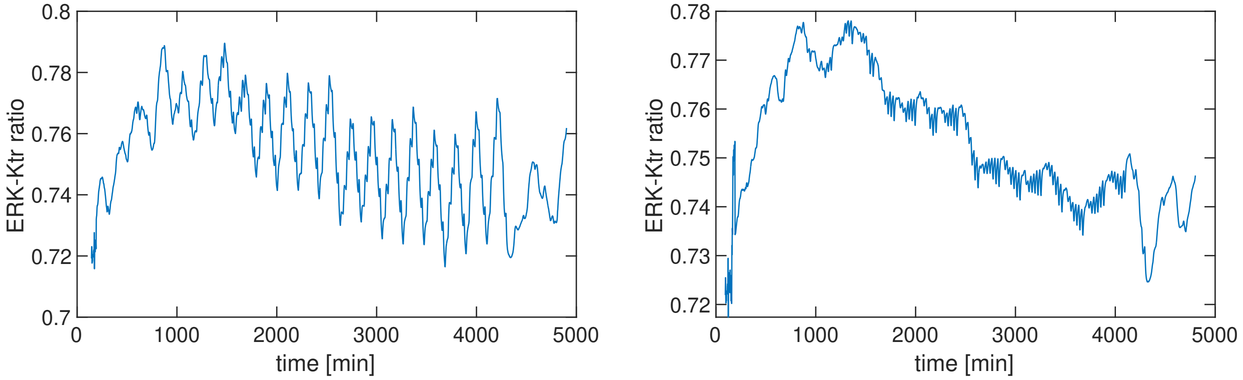


FIG. S9: **Trend of a trajectory**, from detrending with a moving averaging window of 50 frames (left) and 100 frames (right).

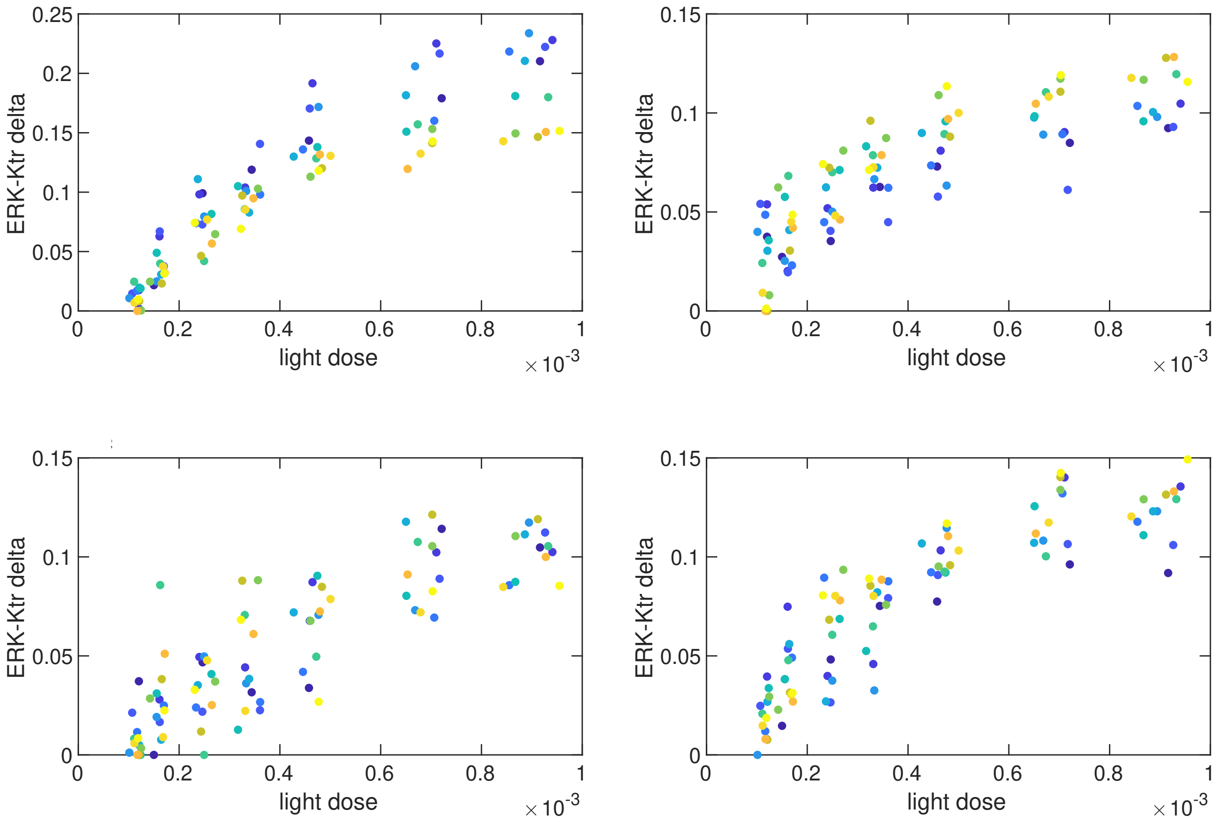


FIG. S10: **Some cells could experience a drift in their encoding scheme.** Dose-responses of 4 different cells from the main dataset (20 repeats) and same stage position (i.e. receiving identical light inputs). The marker color codes for increasing repeat number (blue to yellow). Cells in top panels display opposing drifts in reaction to the light stimuli, while the bottom cells do not display a clear drift. The trajectories are detrended with a moving averaging window of 100 frames (3.3h).

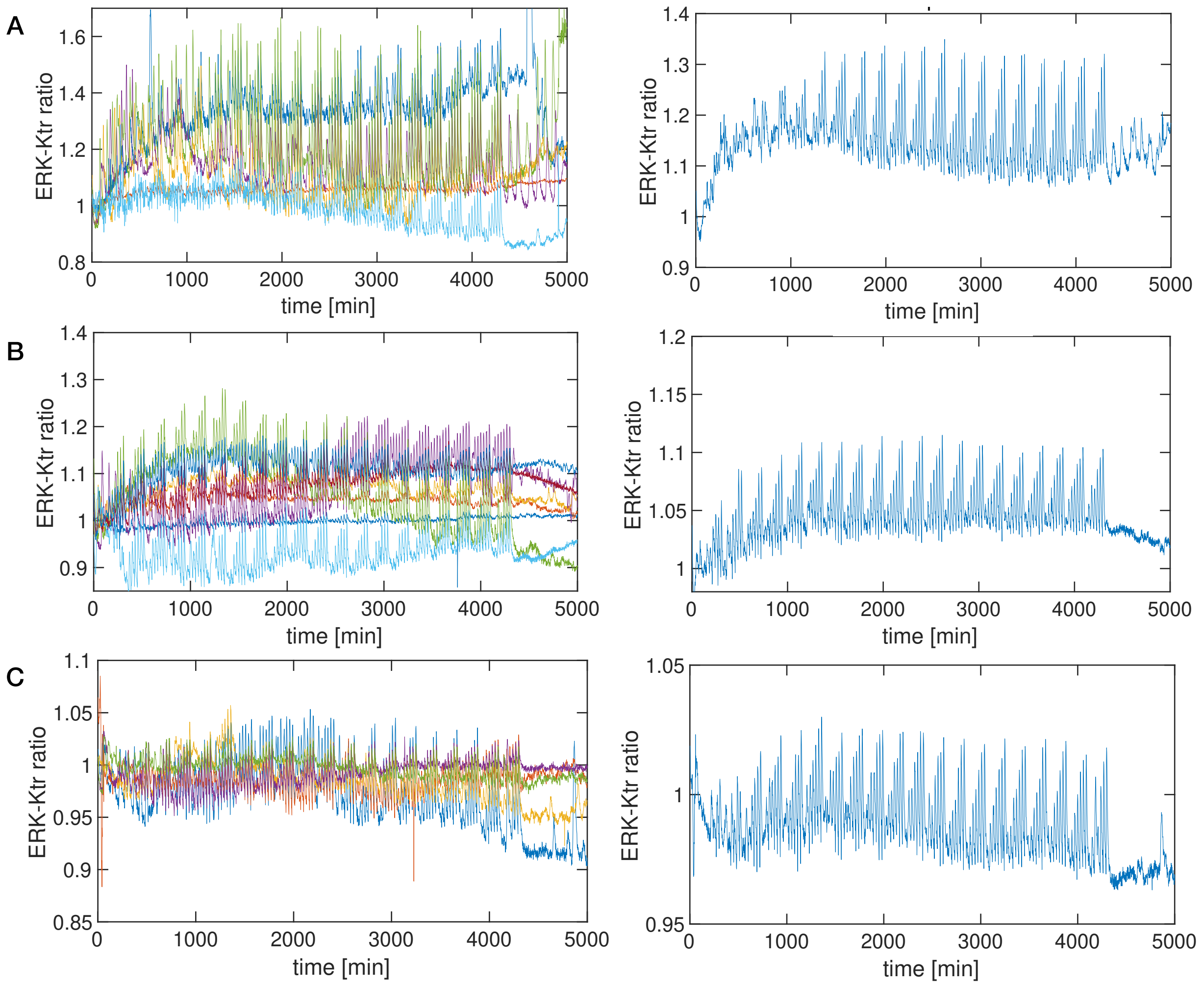


FIG. S11: **Raw ERK-KTR ratio trajectories** (left) and their population average (right) of the main dataset with 20 repeats. Panels (A) to (C) depict cells in three different stage positions that were selected for further analysis.

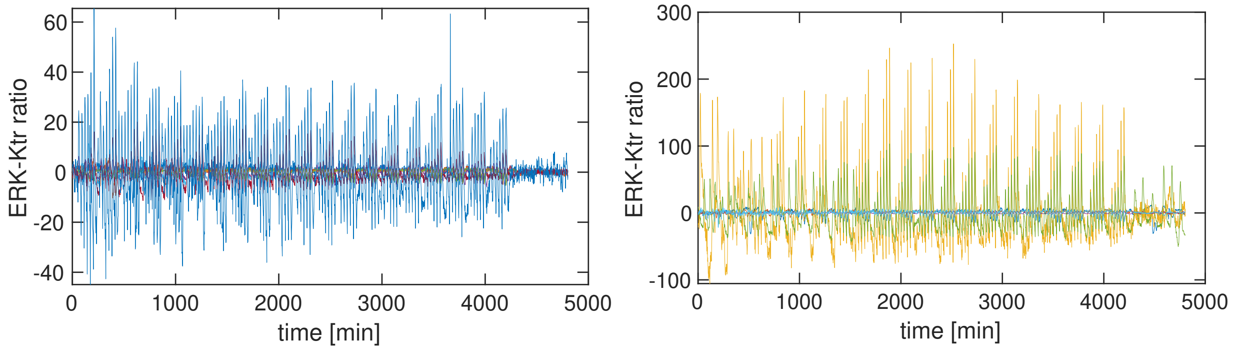


FIG. S12: **Detrended trajectories** from two stage positions of the main dataset with 20 repeats, here detrended with a moving averaging window of 100 frames (3.3h).

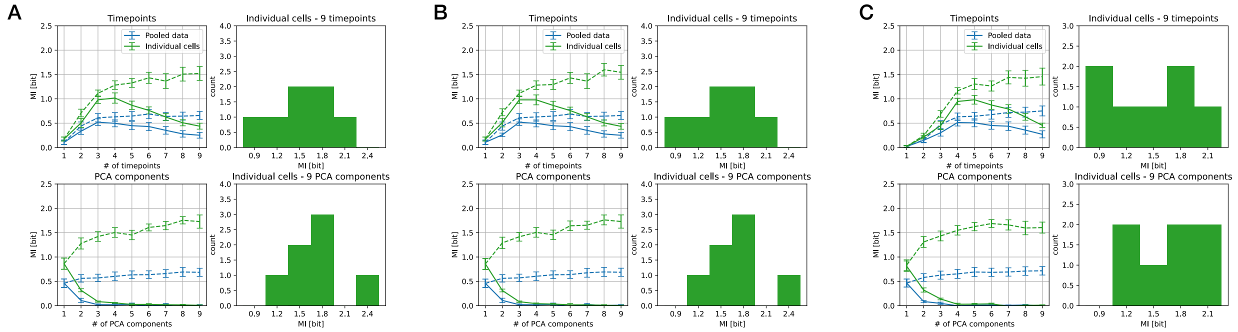


FIG. S13: **The effect of detrending.** Mutual information estimated using raw trajectories as input (A), trajectories detrended using a 4th order polynomial (B) and a moving averaging window of 100 frames (3.3h) (C). Time-point based feature reduction is shown at the top, and PCA at the bottom of each panel. The mutual information estimate shows negligible global differences between the three approaches.

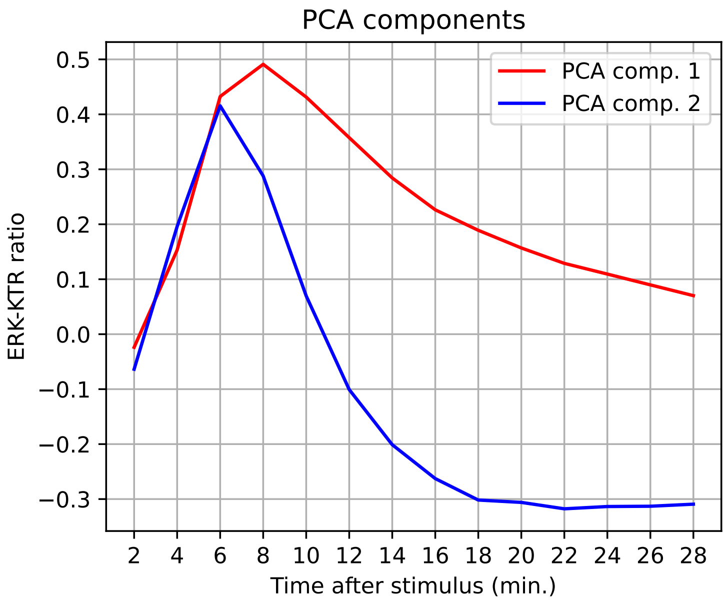


FIG. S14: **Principal component analysis.** First two principal components of the ensemble of the response trajectories from the main dataset with 20 stimulation repeats per condition.

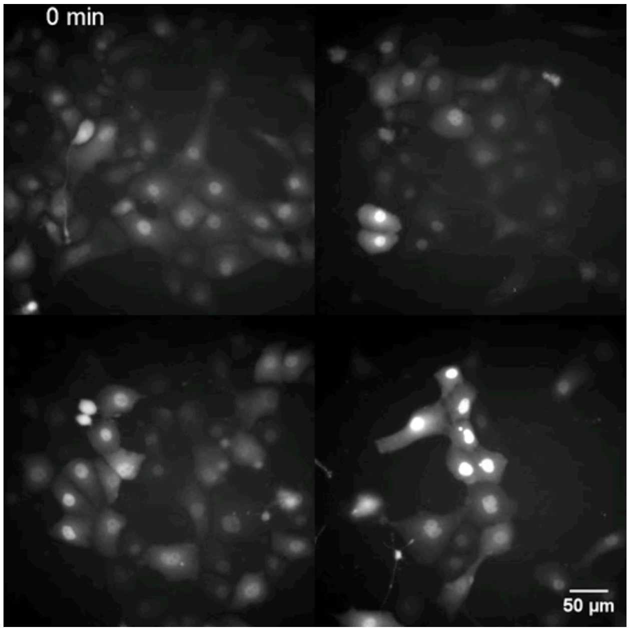


Video 1: **Timelapse of the imaging, first main dataset.** Raw fluorescence microscopy images of the ERK-KTR-mKate2 reporter in cells of the dataset with 20 repeats. The four stage positions are here stitched side by side for easier visualization. An image is taken every 2 min. The application of light stimuli starts at 120 min and ends at 4290 min.

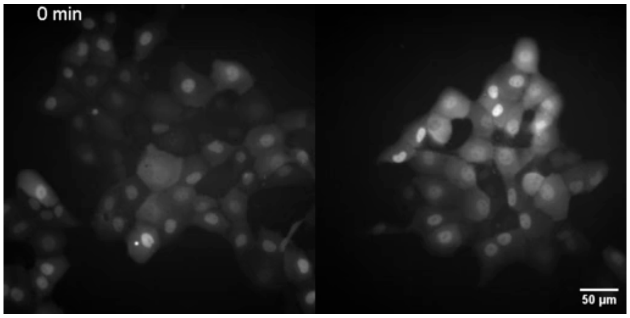


Video 2: **Timelapse of the imaging, second main dataset.** Raw fluorescence microscopy images of the ERK-KTR-mKate2 reporter in cells of the dataset with 15 repeats. The two stage positions are stitched side by side for easier visualization. An image is taken every 2 min. The application of light stimuli starts at 100 min and ends at 3220 min.

[1] L. Hahn, A. M. Walczak, and T. Mora, Dynamical information synergy in biochemical signaling networks, Phys. Rev. Lett. **131**, 128401 (2023).
